# Insight into the genetic network governing long-stalked glandular trichome development in *Nicotiana tabacum*

**DOI:** 10.1101/2025.04.25.650498

**Authors:** Alice Berhin, Aldana Ramirez, Manon Peeters, Maxime Vannieuwenhuyze, Sylvain Legay, Belkacem El Amraoui, Charles Hachez

**Affiliations:** Louvain Institute of Biomolecular Science and Technology, UCLouvain, Louvain-la-Neuve, Belgium; Environmental Research and Innovation, Luxembourg Institute of Science and Technology, Esch-sur-Alzette, Luxembourg

**Keywords:** Nicotiana tabacum, long-stalked glandular trichome, transcriptome, developmental biology, MYB transcription factor, C2H2 zinc finger protein

## Abstract

Glandular trichomes play a pivotal role in plant defense against various biotic and abiotic stresses. To unravel the genetic network driving glandular trichome development in *Nicotiana tabacum*, we employed a transcriptional switch-based approach using heterologous expression of AmMIXTA, a MYB transcription factor from Antirrhinum majus. Through comprehensive RNA-seq analysis of transgenic tobacco seedlings where long-stalked glandular trichome development was induced in aerial parts, we identified a suite of differentially expressed genes, including several transcription factors, shedding light on the early transcriptional cascade governing this developmental process. Moreover, through confocal live cell imaging of transcriptional reporter lines, interaction studies and DAP-seq assays, we confirmed the involvement of *NtZFP8*, a gene identified in our study, in long-stalked glandular trichome development. Our findings support a model in which *NtZFP8* regulates a gene network essential for this developmental process. This study underscores the effectiveness of our approach in decoding the regulatory landscape of glandular trichome development in *N. tabacum* and provides a valuable framework for future functional investigations.

**Key Message:** Glandular trichomes are essential for plant defense. Using a transcriptional switch-based approach in *Nicotiana tabacum*, we identified key regulators, including NtZFP8, which controls long-stalked trichome development. Our study establishes a foundation for future functional analyses of candidate genes uncovered through this approach.

## 1. Introduction

Trichomes, which are epidermal outgrowths found on the aerial parts of many plant species, display an incredible variety of structures, from simple single cells to complex, multicellular forms that are frequently topped with a glandular head (Hauser, 2014). Trichomes play an essential role in plant environmental adaptation by acting as physical barriers against a variety of biotic and abiotic stresses such as excessive sunlight (UV radiation), drought or attacks by pathogens or insects. Furthermore, glandular trichomes produce and store or secrete a diverse array of secondary metabolites - such as acyl sugars, terpenoids, phenylpropanoids, flavonoids, and alkaloids - serving as a chemical line of defense against herbivores and pathogens, and/or to attract pollinators (Gershenzon and Dudareva, 2007, Huchelmann et al., 2017). By enhancing plants resistance to stresses, glandular trichomes contribute to plant survival and reproductive fitness (Hauser, 2014).

In the model plant *Arabidopsis thaliana*, significant progress has been made in understanding the genetic framework governing the development of non-glandular, unicellular branched trichomes, unveiling an extensive network of key regulatory genes. Its genetic basis is managed by a complex of R2R3 MYB/bHLH/WD-repeat transcription factors regulating trichome initiation and morphogenesis: GLABRA1 (AtGL1), GLABRA3 (AtGL3) / ENHANCER OF GLABRA3 (AtEGL3), and TRANSPARENT TESTA GLABRA1 (AtTTG1). For a comprehensive review, refer to Han et al. (2022).

Compared to unicellular trichomes, the genetic basis of multicellular trichome development is less understood but shows ongoing progress (Hauser, 2014, Chalvin et al., 2020). Developmental regulation involves multiple genes and regulatory pathways, varying widely among plant species and trichome types (non-glandular vs glandular, multicellular vs unicellular) and highlights their diversity of evolutionary origin (Hauser, 2014, Chalvin et al., 2020). This complexity hinders the creation of a universal model for understanding the genetic factors in glandular trichome development Unlike *Arabidopsis thaliana*, which provides a unified model for single-cell trichome study, different species or groups serve as models to understand the genetic control behind specific glandular trichome types, such as *Lamiaceae* for peltate trichomes or *Solanaceae* for capitate trichomes (Tissier, 2012).

Our research aims to unravel the transcriptional networks governing the development of these epidermal structures, particularly in *Nicotiana tabacum*, belonging to the agronomically important *Solanaceae* family in the *Asterid* clade. This model plant possesses three multicellular trichome types: short-stalked glandular trichomes, non-glandular trichomes and long-stalked glandular trichomes.

Numerous transcription factors are implicated as regulators of multicellular trichome development across *Asterid* species, indicating the genetic intricacy controlling the development of these structures (Han et al., 2022): (1) MYB transcription factors like AaMIXTA, AaMYB1 in *Artemisia annua* (Shi et al., 2017, Matias-Hernandez et al., 2017), AmMIXTA, AmMYBML1, homologs of AaMIXTA, in *Antirrhinum majus*, SlMX1 in *Solanum lycopersicum* (Ewas et al., 2016, Wu et al., 2023), NbMYB123 in *Nicotiana benthamiana* (Liu et al., 2018), McMIXTA in *Mentha canadensis* (Qi et al., 2022) and GLAND CELL REPRESSOR (SlGCR1 and SlGCR2) (Chang et al., 2024); (2) C2H2 zinc-finger (ZFP) transcription factors such as SlHAIR, SlZFP6, SlZFP8/SlHAIR2 (Chang et al., 2018, Zheng et al., 2021) and GLABROUS INFLORESCENCE STEM (NbGIS) (Liu et al., 2018); (3) bHLH transcription factors such as SlMYC1 (Xu et al., 2018); (4) Homeodomain-leucine zipper (HD-ZIP) IV transcription factors like AaHD1, AaHD8 (Yan et al., 2017, Yan et al., 2018), SlCD2, homolog of AaHD8 (Nadakuduti et al., 2012), SlWOOLLY and NbWOOLLY (Yang et al., 2011a, Yang et al., 2015); (5) WRKY transcription factors such as AaGSW2 (Xie et al., 2021); (6) AP2/ERF transcription factors like LEAFLESS (SlLFS) and SlTOE1B (Wu et al., 2023, Chang et al., 2024); (7) WUSCHEL-related homeobox such as SlWOX3 (Wu et al., 2023); (8) TOESINTE BRANCHED, CYCLOIDEA, and PROLIFERATING CELL NUCLEAR ANTIGEN BINDING FACTOR transcription factor (TCP) like BRANCHED2a (SlBRC2a) (Wu et al., 2024). Additionally, cell cycle regulators, like E3 ubiquitin ligase (SlMTR1/SlCYCB2 and SlMTR2, NbCYCB2 and NtCYCB2) impact trichome development, emphasizing the sophisticated control of cell division and differentiation needed for multicellular trichome architecture in *Solanaceae* (Wu et al., 2020, Gao et al., 2017, Wang et al., 2021, Wu et al., 2023).

Recent studies in tomato underscore the pivotal role of HD-ZIP transcription factors in determining trichome cell fate, revealing a regulatory landscape distinct from that of unicellular trichome development (Wu et al., 2023, Wu et al., 2024). Indeed, SlWOOLLY controls trichome development in a dosage-dependent mechanism controlled by self-activating and SlMTR1-mediated feedback. Trichome type fate is then controlled by SlWOX3 and SlMX1 for digitate trichomes and SlLFS for peltate trichomes; high levels of SlWOOLLY preferentially activate SlWOX3/SlMX1 complex, which inhibits *SlLFS* expression (Wu et al., 2023). When *SlWOOLLY* expression level is low, SlBRC2a mediates a negative feedback loop, transitioning cells from division to expansion (Wu et al., 2024). In addition, SlGCR1/SlGCR2 operate dose-dependently in the apical cell to regulate glandular head formation (Chang et al., 2024); SlTOE1B inhibits *SlGCRs* expression whose low expression level activates *SlLFS* leading to gland formation (Chang et al., 2024).

In *N. tabacum*, glandular trichome development is regulated through a transcriptional pathway differing from the regulatory mechanisms controlling the formation of unicellular non-glandular trichomes. (Glover et al., 1998, Payne et al., 1999, Perez-Rodriguez et al., 2005). In *A. majus*, AmMIXTA regulates cell wall deposition in floral papillae (Noda et al., 1994). Ectopic expression of *AmMIXTA* induces the formation of multicellular trichomes in *N. tabacum*, demonstrating the involvement of R2R3 MYB transcription factors in the development of glandular trichomes (Payne et al., 1999, Perez-Rodriguez et al., 2005, Glover et al., 1998).

By using inducible *AmMIXTA* expression as a switch to trigger long-stalked glandular trichome formation in a normally trichome-free organ such as the cotyledon, this exploratory study opens new avenues for investigating the molecular mechanisms that initiate and shape early genetic events in glandular trichome development in tobacco. Our work provides a broader framework for understanding the initiation of glandular trichome development in *N. tabacum* and identifies potential candidate genes for future research in this field.

## 2. Results

### 2.1 Ectopic Induction of *AmMIXTA* Unravels Genes Integral to Long-stalked Glandular Trichome Development

In this study, we investigated the transcriptional cascade triggered by *AmMIXTA* heterologous expression and leading to the induction of supernumerary long-stalked glandular trichomes in *N. tabacum* (Payne et al., 1999, Perez-Rodriguez et al., 2005, Glover et al., 1998), induction of short-stalked glandular or non-glandular trichome development was not observed. Driving *AmMIXTA* expression with a β-estradiol inducible promoter (Curtis and Grossniklaus, 2003), we aimed to dissect the molecular mechanisms governing long-stalked glandular trichome initiation and development on the cotyledons of tobacco.

Expression of *AmMIXTA* resulted in a pronounced trichome-rich phenotype on the cotyledons and hypocotyl, as evidenced by comparative analysis of induced and non-induced plants (Figure 1A-E and Figure S1A-B). Microscopic examination revealed an increase in long-stalked glandular trichome density from 1.67±5.34 cm^-2^ to 664.33±346.58 cm^-2^ upon *AmMIXTA* induction (Figure 1E), confirming its role in redirecting protodermal cell fate towards trichome formation (Payne et al., 1999, Perez-Rodriguez et al., 2005, Glover et al., 1998). Ectopically induced long-stalked trichomes exhibited a morphology similar to naturally occurring ones, with chlorophyll autofluorescence indicating the presence of chloroplasts in glandular cells (Figure S1C-D). In addition, rhodamine B staining, a dye used for acylsugar labeling, marked the trichome heads, suggesting secondary metabolite biosynthesis activities in the gland (Figure S1E). Nuclear localization of AmMIXTA was confirmed using a VENUS-tagged version of the protein (Figure S1F).

**Figure 1:**
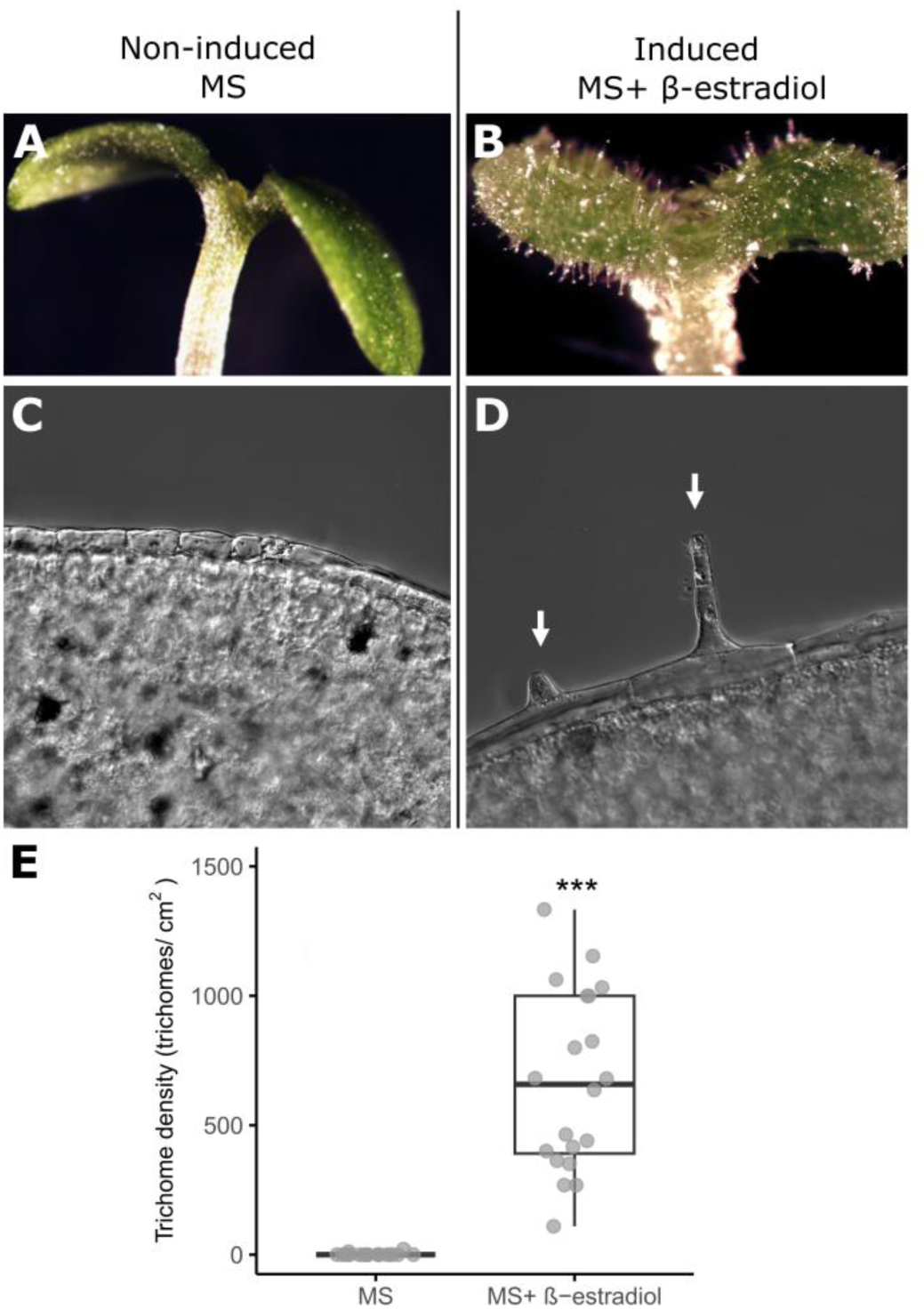
Phenotype of pEST::AmMIXTA transgenic plants. (A-D) Ten-day-old pEST::AmMIXTA seedlings germinated on non-inducing medium (1⁄2 MS medium) (A/C) and inducing medium (1⁄2 MS medium supplemented with 10 μM β-estradiol) (B/D). (C/D) Differential Interference Contrast (DIC) pictures of the epidermal layer. Arrows indicate developing trichomes. Images were captured at 40X magnification. (E) Trichome density on induced and non-induced seedlings presented as a boxplot (n=20 plants). Individual data points are depicted as dots. Asterisks denote significant differences from the MS condition as determined by Student’s t-test: ***p < 0.001

Using the *AmMIXTA* inducible system, we conducted RNA-seq to identify genes crucial for initiating and driving the development of long-stalked glandular trichomes. Seedlings treated with 10 μM β-estradiol for 4 h and 48 h were compared to non-induced controls (Figure S2). RT-qPCR validation confirmed significant upregulation of *AmMIXTA* transcripts, by 37-fold at 4 h and 138-fold at 48 h (Figure 2A), with trichome formation observed prominently at 48 h. Subsequent RNA-seq analysis yielded a high-quality mapping to the *N. tabacum* genome (Sierro et al., 2014) (Table S1), identifying 719 and 1947 differentially expressed genes (DEGs) at the 4 h and 48 h time points (0.67>Fold change and Fold change> 1.5, p-value <0.05) (Table S2A and S2B). Emphasis was placed on analyzing upregulated genes in subsequent data analyses due to this cell fate reprogramming. Among these, 166 and 604 genes were upregulated at 4 h and 48 h, respectively, with some genes upregulated at both time points (Fold change >1.5, p-value <0.05) (Figure 2B and Table S2A). The 4 h post-*AmMIXTA* induction captures genes involved in early stages of long-stalked glandular trichome development, while 48 h post-induction captures cell trans-differentiation into glandular trichome cells. Notably, four genes including transcription factors were consistently upregulated at both time points: two bHLHs, one AP2/ERF, one TRIHELIX transcription factors and one protein containing a MYB domain, (Figure 2B-C).

**Figure 2:**
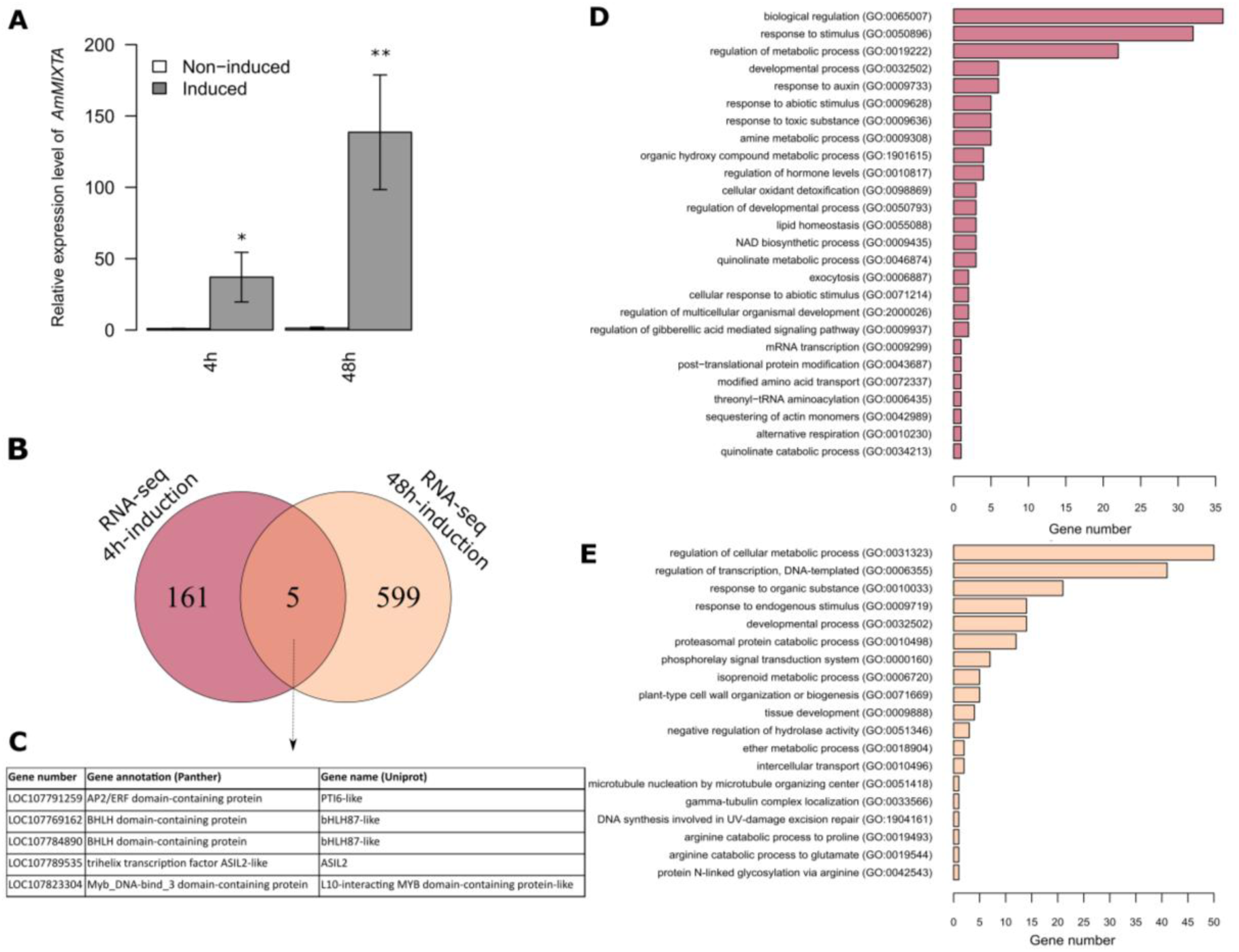
RNA-seq Analysis 4 h and 48 h After β-Estradiol Induction of pEST::AmMIXTA. (A) Expression level of AmMIXTA in 10-day-old seedlings germinated on 1/2 MS medium and subjected to β-estradiol induction for 4 h and 48 h compared to the control in non-inducible conditions (1/2 MS). Expression normalization was performed using the geometric averaging of multiple housekeeping genes (NtEF2, NtACTIN, NtUBC2). Values represent means ± SE, n=3. Asterisks denote significant differences compared to non-induced conditions as determined by Student’s t-test: *p < 0.05; **p < 0.001. (B) Differentially up-regulated genes between the pEST::AmMIXTA seedlings induced with β-estradiol for 4 hours and 48 hours, and their own non-induced controls, represented by a Venn diagram. (C) Differentially up-regulated genes common to both analyses. (D/E) GO functional enrichment analysis based on the differentially up-regulated genes at 4 h (D) and 48 h (E). The graphs represent categories with a statistically significant difference between induced and non-induced plants (p-value < 0.05). The x-axis represents the fold enrichment of genes observed in the RNA-seq data over the expected number in the reference list. The y-axis shows the over-represented GO terms from the Biological Process classification according to PANTHER.

Gene Ontology (GO) term enrichment analysis via PANTHER (p-value < 0.05) illustrates distinct biological function enrichment patterns of upregulated DEGs at 4 h and 48 h post-*AmMIXTA* induction, reflecting the dynamic nature of trichome development (Figure 2D-E; Figure S3, Table S3) (Mi et al., 2019, Thomas et al., 2022). At 4 hours post-induction, enrichment included terms associated among others with developmental processes and responses to stimuli (Figure 2D, Table S3), and at 48 hours with developmental processes (Figure 2E, Table S3) and secondary metabolite biosynthesis (including isoprenoid metabolic process). The observed enrichments align with expected GO categories for glandular trichome development and differentiation.

### 2.2 Transcription Factors Involved in Long-stalked Glandular Trichome Initiation and Development

Further dissecting the dataset, we cross-referenced the upregulated genes at both time points with genes referenced as transcription factor in the Plant TFBD (Jin et al., 2017) database and on PANTHER (Thomas et al., 2022). This yielded two distinct lists: one comprising 15 transcription factors at 4 hours, likely involved in the initiation of long-stalked glandular trichome development, and another consisting of 32 candidates at 48 hours, associated with further development and differentiation (Table 1).

**Table 1:**
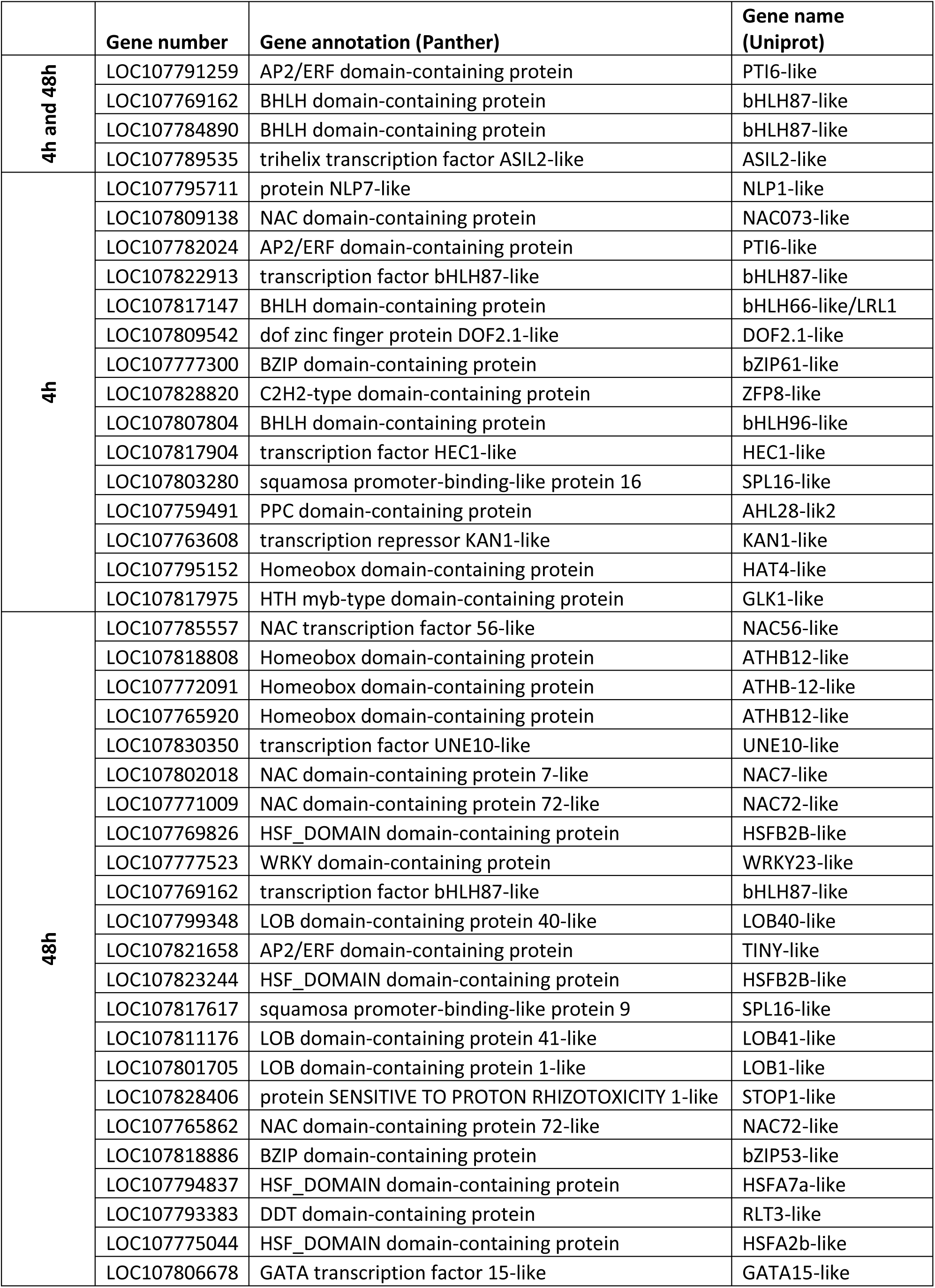
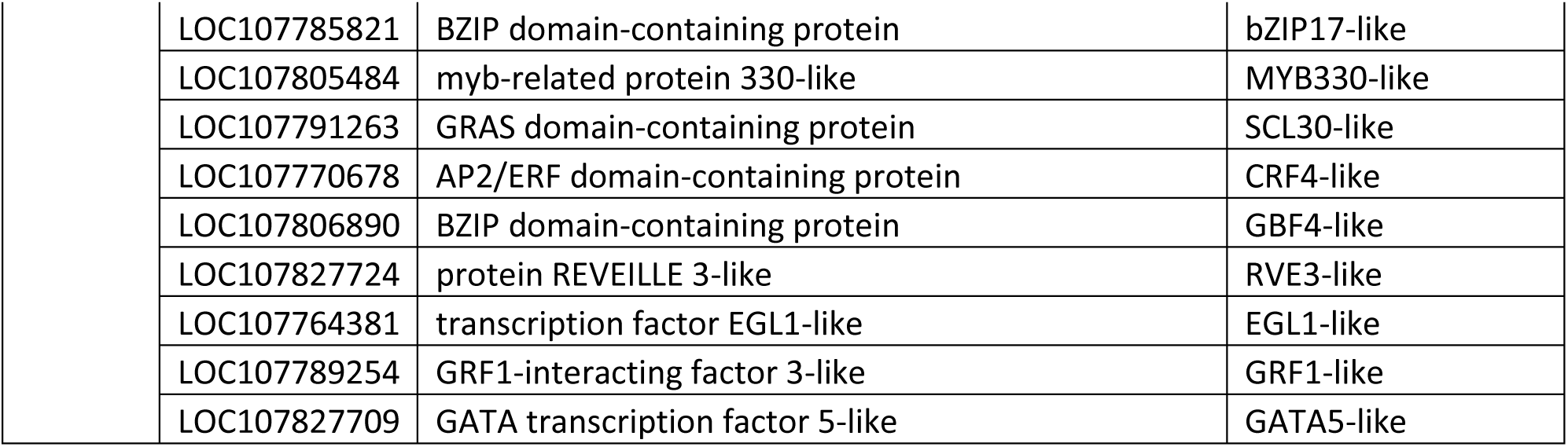
Transcription Factors Induced at 4 h and/or 48 h. This table lists transcription factors that were induced at 4 hours and 48 hours post-induction,

|  | Gene number | Gene annotation (Panther) | Gene name (Uniprot) |
| --- | --- | --- | --- |
| 4h and 48h | LOC107791259 | AP2/ERF domain-containing protein | PTI6-like |
|  | LOC107769162 | BHLH domain-containing protein | bHLH87-like |
|  | LOC107784890 | BHLH domain-containing protein | bHLH87-like |
|  | LOC107789535 | trihelix transcription factor ASIL2-like | ASIL2-like |
| 4h | LOC107795711 | protein NLP7-like | NLP1-like |
|  | LOC107809138 | NAC domain-containing protein | NAC073-like |
|  | LOC107782024 | AP2/ERF domain-containing protein | PTI6-like |
|  | LOC107822913 | transcription factor bHLH87-like | bHLH87-like |
|  | LOC107817147 | BHLH domain-containing protein | bHLH66-like/LRL1 |
|  | LOC107809542 | dof zinc finger protein DOF2.1-like | DOF2.1-like |
|  | LOC107777300 | BZIP domain-containing protein | bZIP61-like |
|  | LOC107828820 | C2H2-type domain-containing protein | ZFP8-like |
|  | LOC107807804 | BHLH domain-containing protein | bHLH96-like |
|  | LOC107817904 | transcription factor HEC1-like | HEC1-like |
|  | LOC107803280 | squamosa promoter-binding-like protein 16 | SPL16-like |
|  | LOC107759491 | PPC domain-containing protein | AHL28-lik2 |
|  | LOC107763608 | transcription repressor KAN1-like | KAN1-like |
|  | LOC107795152 | Homeobox domain-containing protein | HAT4-like |
|  | LOC107817975 | HTH myb-type domain-containing protein | GLK1-like |
| 48h | LOC107785557 | NAC transcription factor 56-like | NAC56-like |
|  | LOC107818808 | Homeobox domain-containing protein | ATHB12-like |
|  | LOC107772091 | Homeobox domain-containing protein | ATHB-12-like |
|  | LOC107765920 | Homeobox domain-containing protein | ATHB12-like |
|  | LOC107830350 | transcription factor UNE10-like | UNE10-like |
|  | LOC107802018 | NAC domain-containing protein 7-like | NAC7-like |
|  | LOC107771009 | NAC domain-containing protein 72-like | NAC72-like |
|  | LOC107769826 | HSF_DOMAIN domain-containing protein | HSFB2B-like |
|  | LOC107777523 | WRKY domain-containing protein | WRKY23-like |
|  | LOC107769162 | transcription factor bHLH87-like | bHLH87-like |
|  | LOC107799348 | LOB domain-containing protein 40-like | LOB40-like |
|  | LOC107821658 | AP2/ERF domain-containing protein | TINY-like |
|  | LOC107823244 | HSF_DOMAIN domain-containing protein | HSFB2B-like |
|  | LOC107817617 | squamosa promoter-binding-like protein 9 | SPL16-like |
|  | LOC107811176 | LOB domain-containing protein 41-like | LOB41-like |
|  | LOC107801705 | LOB domain-containing protein 1-like | LOB1-like |
|  | LOC107828406 | protein SENSITIVE TO PROTON RHIZOTOXICITY 1-like | STOP1-like |
|  | LOC107765862 | NAC domain-containing protein 72-like | NAC72-like |
|  | LOC107818886 | BZIP domain-containing protein | bZIP53-like |
|  | LOC107794837 | HSF_DOMAIN domain-containing protein | HSFA7a-like |
|  | LOC107793383 | DDT domain-containing protein | RLT3-like |
|  | LOC107775044 | HSF_DOMAIN domain-containing protein | HSFA2b-like |
|  | LOC107806678 | GATA transcription factor 15-like | GATA15-like |

**Table 1: Transcription Factors Induced at 4 h and/or 48 h.**
|  |  |  |
| --- | --- | --- |
| LOC107785821 | BZIP domain-containing protein | bZIP17-like |
| LOC107805484 | myb-related protein 330-like | MYB330-like |
| LOC107791263 | GRAS domain-containing protein | SCL30-like |
| LOC107770678 | AP2/ERF domain-containing protein | CRF4-like |
| LOC107806890 | BZIP domain-containing protein | GBF4-like |
| LOC107827724 | protein REVEILLE 3-like | RVE3-like |
| LOC107764381 | transcription factor EGL1-like | EGL1-like |
| LOC107789254 | GRF1-interacting factor 3-like | GRF1-like |
| LOC107827709 | GATA transcription factor 5-like | GATA5-like |

Both lists featured key transcription factor families known for their role in trichome development in *Solanaceae*, including bHLHs, C2H2 zinc-finger proteins, MYBs, HD-ZIPs and WRKYs. This approach highlighted the transcription factors critical for both the onset and progression of long-stalked glandular trichome development, illustrating the complexity of the regulatory network involved.

### 2.3 NtZFP8 is Expressed in Developing Trichomes

Various C2H2 zinc finger protein transcription factors, including AtZFP6, AtGIS, AtGIS3, NbGIS, JcZFP8, SlHAIR and SlHAIR2, play significant roles in trichome formation in *Arabidopsis*, tomato, and tobacco (Gan et al., 2007, Zhou et al., 2013, Sun et al., 2015, Liu et al., 2018, Shi et al., 2018, Zheng et al., 2021).

Through our analysis, NtZFP8 (LOC107828820) emerged as a noteworthy ZFP transcription factor, exhibiting significant upregulation 4 hours following the induction of *AmMIXTA*. To understand its role in trichome development in *Nicotiana tabacum*, we examined its expression across various tissues. RT-qPCR results showed that *NtZFP8* expression is higher in developing leaves than in mature leaves, suggesting a role during trichome initiation and development (Figure 3B). This was further confirmed using a NtZFP8 promoter reporter line (*pNtZFP8:nlsGFP-GUS*), which demonstrated promoter activity localized to the epidermal layer during early developmental stages (Figure 3A). In leaf primordia, NtZFP8 promoter activity was observed in epidermal pavement cells, stomata, non-glandular trichomes, long-stalked glandular trichomes (stalk and developing glandular heads at 1-2 cell stages), and short-stalked glandular trichomes (stalk and head). This expression is consistently present in developing trichomes but ceases post-development. In mature leaves, *NtZFP8* expression is no longer detectable in trichomes but remains active in some epidermal pavement cells and stomata (Figure S4B).

**Figure 3:**
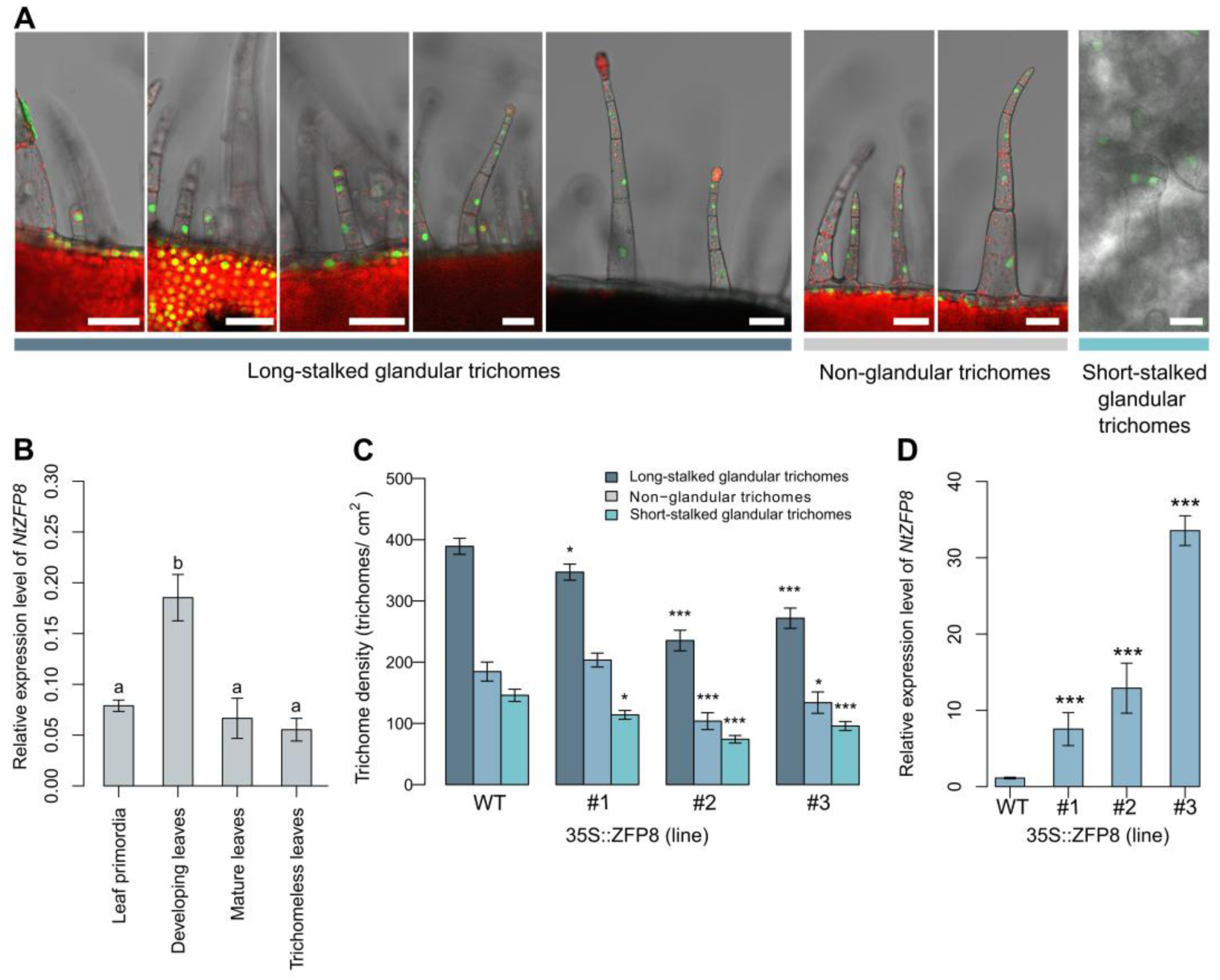
NtZFP8 Expression Pattern in leaves. (A) Reporter lines pZFP8::nlsGFP-GUS showing expression in different types of trichomes during their development in expanding leaves of 6-week-old plants. Scale bars represent 50 μm; green indicates nlsGFP; red denotes chlorophyll autofluorescence. (B) RT-qPCR analysis of different tissues (leaf primordia, developing leaves, mature leaves, and leaves without trichomes) from WT plants. Gene expression levels are expressed relative to the geometric mean of three different internal controls (NtEF2, NtACTIN, NtUBC2). Results are displayed as mean ± SE, n=4. Significant differences were determined by ANOVA, with each letter indicating a significant variation between tissues (p < 0.05) (C) Trichome density in different independent lines p35S::ZFP8. (D) Expression level of *NtZFP8* in the overexpressing lines measuring by RT-qPCR. Gene expression levels are expressed relative to the WT and the geometric mean of three different internal controls (*NtEF2, NtACTIN, NtUBC2*). Asterisks denote significant differences to WT as determined by Student’s t test: ***p < 0.001, **p<0.01 and *p<0.05.

In *NtZFP8* overexpressing lines (*p35S::NtZFP8*), the density of all three trichome types in fully grown leaves was generally significantly reduced compared to WT plants (Figure 3C-D). This phenotype suggests that NtZFP8 may influence trichome development in *N. tabacum*.

### 2.4 Convergent Transcriptional Targets of AmMIXTA and NtZFP8 in Tobacco

We investigated a potential physical interaction between AmMIXTA and NtZFP8 using a yeast-2-hybrid assay (Paiano et al., 2019). *AmMIXTA* coding sequence (CDS) was fused to the GAL4 activation domain (AD), and *NtZFP8* CDS to the GAL4 binding domain (BD). Colonies containing AD-AmMIXTA and BD-NtZFP8 grew in a similar way on SD-WL (selective medium without tryptophan and leucine). A significant growth differences on SD-HAWL (medium without of histidine and adenine) was observed and indicated a physical interaction between AmMIXTA and NtZFP8 (Figure 4).

**Figure 4:**
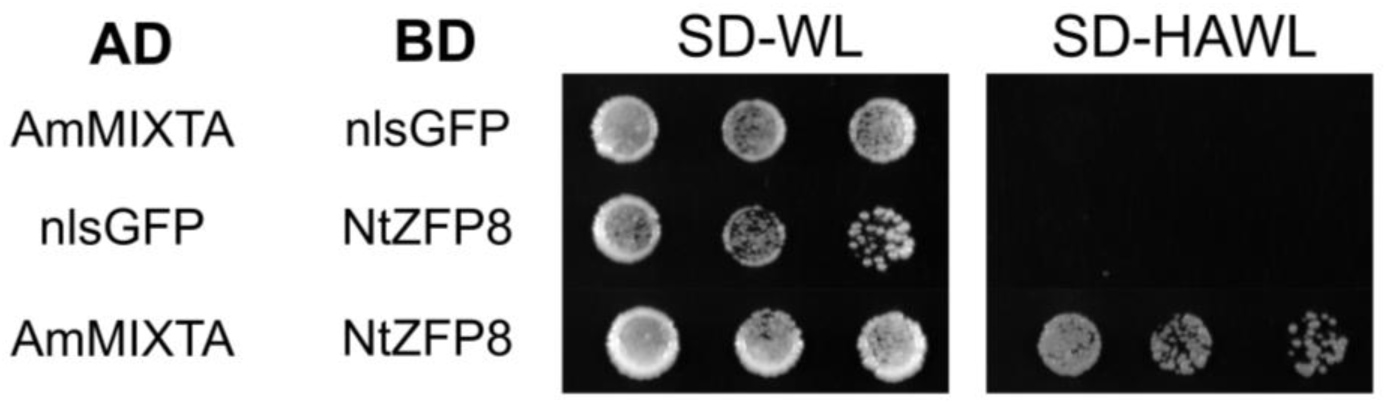
Interaction Between NtZFP8 and AmMIXTA Demonstrated by Yeast Two-Hybrid Assay. This assay was performed using a control medium lacking tryptophan and leucine (SD/-WL) and a selective medium devoid of tryptophan, leucine, histidine, and adenine (SD/-HAWL) to investigate the interaction between AmMIXTA and NtZFP8. NlsGFP served as the negative control for the interaction. ’AD’ represents the activation domain, and ’BD’ denotes the binding domain; ’SD’ refers to synthetically defined medium. The experiment was replicated three times, consistently showing similar results.

In order to unravel downstream pathways regulated by AmMIXTA and NtZFP8 and so identify the transcriptional targets of both transcription factors, we employed DNA affinity purification sequencing (DAP-seq) (Bartlett et al., 2017). DNA sequences that were bound to the transcription factors were sequenced and mapped to the *N. tabacum* genome (Sierro et al., 2014) (Table S4). Hits were identified by comparing against the negative control (nlsGFP protein), (peak shape score <2 and p-value < 0.05) to delineate significant binding regions.

Among all the regions significantly enriched in the analysis, 6.4% (1874 hits) for AmMIXTA and 5.8% (1235 hits) for NtZFP8 were in regions surrounding annotated genes (-3kb and 1kb respectively before the start codon and after the stop) corresponding respectively to 1162 and 974 distinct genes. The rest were in intergenic regions and were not further included in the subsequent analysis (Table S5 and S6). Among the genes of interest, 425 genes targeted by AmMIXTA and 265 by NtZFP8 were annotated as uncharacterized. Analysis of the binding site locations surrounding annotated genes revealed that, for both AmMIXTA and NtZFP8, a significant majority (over 80%) of the interactions were localized to regions upstream of the START codon, encompassing promoter regions and 5’ untranslated regions (UTRs), as well as within introns (Figure 5A-B).

**Figure 5:**
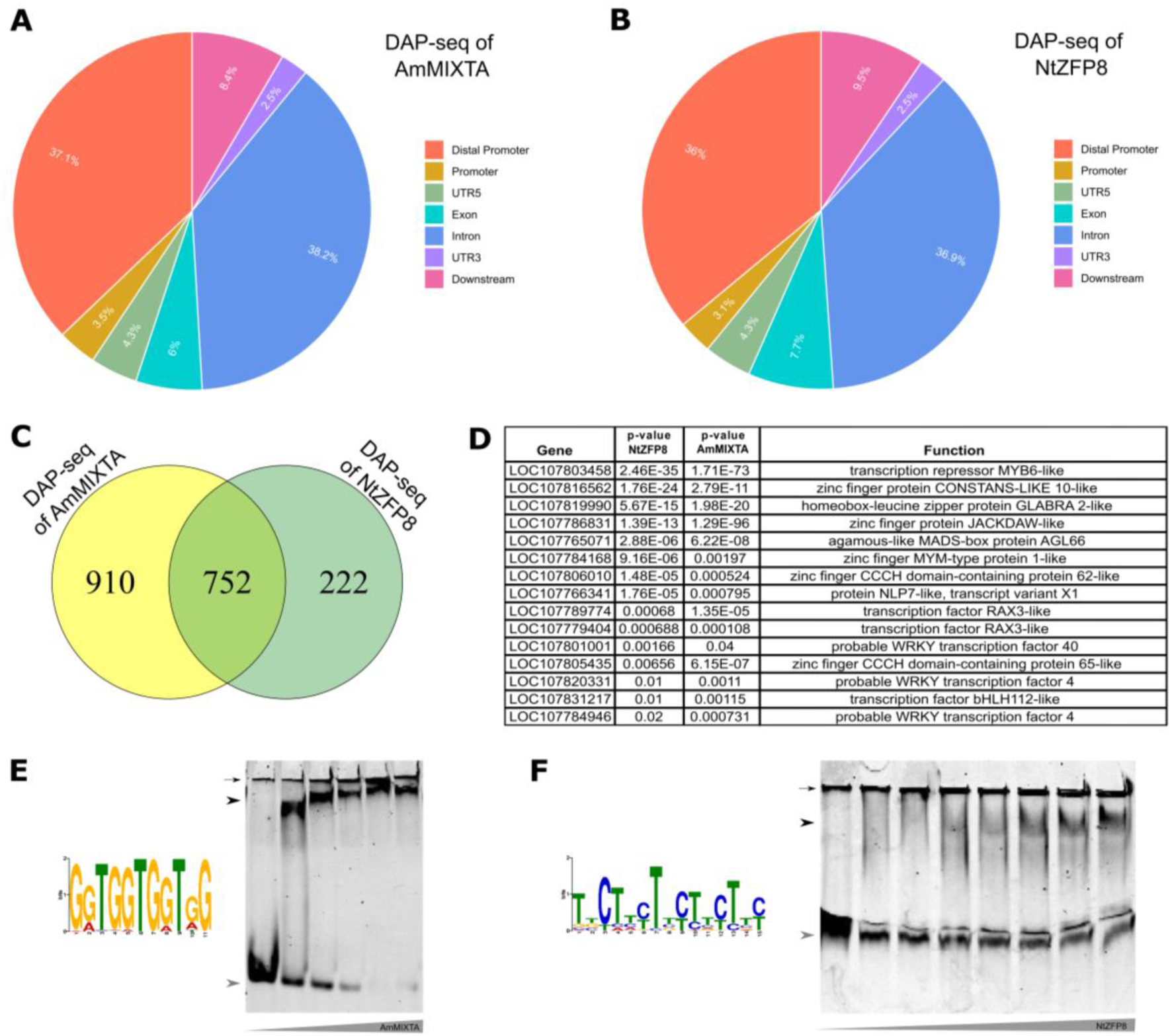
Direct DNA Targets of AmMIXTA and NtZFP8 Transcription Factors. Distribution of (A) AmMIXTA-binding peaks relative to gene structures and (B) NtZFP8-binding peaks. Definitions for gene regions: Distal Promoter (from -3kb to -0.5kb before the START codon); Promoter (from -0.5kb up to the 5’ UTR); UTR5: 5’ Untranslated region, UTR3: 3’ Untranslated region, Exon, Intron and Downstream Region (from the end of the 3’ UTR to 1kb after the STOP codon). (C) Venn diagram showing the overlap between AmMIXTA and NtZFP8 targets. (D) Identification of the top 15 transcription factors shared by AmMIXTA and NtZFP8, as revealed through the DAP-seq assay. (E) The top-scoring motif, GGTGGTGGTGG, was identified with an e-value of 2.2x10^−902^, according to MEME analysis. Gel shift assay shows a binding of AmMIXTA to the fluorescent DNA substrate: GGTGGTGGTGG; Protein concentration were used from 0 to 3000nM. (F) The top-scoring motif, TBCTTYTYCTTCTTY was identified with an e-value of 2.6x10^−76^ according to MEME analysis. Gel shift assays shows a binding of NtZFP8 to the fluorescent DNA substrate: CTTCTTCTTCT; Protein concentration were used from 0 to 4400nM. (B). black arrow head, bound DNA; Grey arrow head, unbound DNA; Small black arrow, wells.

Utilizing the MEME suite for motif discovery (Bailey et al., 2015), we identified specific DNA sequences preferentially bound by each transcription factor: AmMIXTA showed a strong affinity for the GGTGGTGGTGG motif, with an e-value of 2.2x10^−902^ (Figure 5E), whereas NtZFP8 predominantly bound to the TBCTTYTYCTTCTTY motif, with an e-value of 2.6x10^−76^ (Figure 5F). Motif binding sites were confirmed using electrophoretic mobility shift assay (EMSA) confirming a binding to CTTCTTCTTCT for NtZFP8 and to GGTGGTGGTGG for AmMIXTA, a motif similar to the GGTAGGT motif already identified for several MYB proteins (JASPAR database) (Rauluseviciute et al., 2024) (Figure 5E-F).

When examining the overlap in genes targeted by both transcription factors, we identified 752 common genes, representing 45% and 77% of the total number of genes targeted by AmMIXTA and NtZFP8, respectively (Figure 5C). Notably, the top 15 shared targets of both proteins primarily binding in regions upstream of the START codon, mainly comprised genes encoding key transcription factor families such as MYBs, zinc fingers, bHLHs, HD-ZIPs, and WRKYs, indicating a focused regulatory impact on essential components of the plant transcriptional machinery known to impact epidermal patterning, (Figure 5D and Figure S5).

## 3. Discussion

Our research exploited ectopic expression of *AmMIXTA*, acting as a transcriptional switch inducing long-stalked glandular trichome formation to elucidate the transcriptional cascade leading to the development of those trichomes in *N. tabacum*.

Results confirmed AmMIXTA as a master regulator orchestrating a complex developmental cascade, specifically inducing long-stalked glandular trichomes in cotyledon epidermal cells without affecting other cell types (Figure 1 and 2A and Figure S1) (Glover et al., 1998, Payne et al., 1999). This suggests the presence of an endogenous MIXTA-like homolog in tobacco, likely regulating glandular trichome development through a transcriptional network similar to the one described in this study.

### 3.1 Transcription Factors at the Crossroads of Trichome Initiation and Development

Leveraging RNA-seq analysis at critical post-induction intervals (4 h and 48 h) allowed us to capture the dynamic nature of the transcriptional response elicited by AmMIXTA. Focusing on upregulated genes, the significant increase in differentially expressed genes (DEGs) from 166 at 4 hours to 604 at 48 hours not only illustrates the expanding regulatory influence of AmMIXTA over time but also suggests a phased activation of the downstream transcriptional network (Figure 2 A-C and Figure S2).

Our research examines transcriptional regulation of trichome development, focusing on transcription factors known for their roles in trichome development across various plant species (Table 1). Although a role in cuticle and cell wall formation can be hypothesized for AmMIXTA as is the case of its AaMIXTA homolog (Noda et al., 1994, Shi et al., 2017), several of the identified transcription factors or their close homologs have already been associated with trichome development in previous studies.

MYB transcription factors, including AtMYB106 and AtGL1, are essential for trichome initiation and development (Han et al., 2022). We identified NtGOLDEN2-like protein (GLK1) and NtKANADI (KAN1) at the trichome initiation stage (4 h post-induction). In Arabidopsis, GOLDEN2-like proteins support cellular differentiation (Chen et al., 2016), while AtKAN1 regulates trichome development by interacting with AtGL1 (Wang et al., 2019). HD-ZIP IV members, such as SlWOOLLY and AtGL2, and HD-ZIP I members, like CsGL1, TINY BRANCHED HAIR (CsTBH), and MICRO-TRICHOME (CsMICT), are crucial for trichome morphogenesis (Khosla et al., 2014, Wu et al., 2023, Wu et al., 2024, Han et al., 2022). Our study identified NtATHB12 (HD-ZIP I) as a potential contributor to trichome development. Our RNA-seq analysis also revealed several bZIP factors potentially influencing trichome development in *N. tabacum*, including *NtbZIP53*, which is connected to jasmonate responses in *Arabidopsis* (Wu et al., 2022). The importance of bHLH transcription factors in trichome development is well documented: AtGL3, AtEGL3 (Payne et al., 2000, Zhang et al., 2003) and SlbHLH95 (Chen et al., 2020). Here, we identified *NtEGL1* (LOC107764381) as upregulated after 48 hours. Its closest Arabidopsis ortholog is *AtGL3*, gene coding for a well-characterized regulator of trichome development. Another bHLH gene, *NtbHLH066* (LOC107817147), was upregulated at 4 hours. It is closely related to *AtbHLH066/Lj-RHL1-LIKE1* (AtLRL1), a gene linked to root hair elongation and branching. In Arabidopsis, AtLRL1 function is suppressed by AtGL2 in non-hair cells, suggesting a possible role in trichome development (Bruex et al., 2012, Lin et al., 2015).

WRKY transcription factors, such as NtWRKY23, may also play roles in trichome development and differentiation (Ishida et al., 2007, Grunewald et al., 2012, Xie et al., 2021). C2H2-type zinc finger proteins are important in developmental processes and stress responses (Chang et al., 2018, Liu et al., 2018, Zheng et al., 2021, Gao et al., 2021), regulating trichome development through hormonal pathways (Liu et al., 2024). In our study, NtZFP8 was significantly upregulated at 4 hours, suggesting its involvement in trichome development.

In the current model describing the transcriptional network regulating glandular trichome development in *S. lycopersicum*, SlWOOLLY is needed to control the balance between peltate and digitate trichome development via SlMX1, among others (Wu et al., 2024). Here, NtWOOLLY was not found to be induced by the overexpression of *AmMIXTA*, suggesting that AmMIXTA could be acting downstream or even independently from this latter. In our case *AmMIXTA* is sufficient to induce long-stalked glandular trichome formation without impacting non-glandular and short-stalked trichome development. This could be related to the activity of SlMX1, AmMIXTA closest homolog in tomato (Wu et al., 2024). SlMX1 has been shown to work alongside SlWOX3 to increase the number of digitate trichomes while reducing the number of peltate trichomes (Ewas et al., 2016).

### 3.2 Regulation of Trichome Development by NtZFP8 and Its Interaction with AmMIXTA

*NtZFP8* is a close ortholog of the uncharacterized *SlZFP8* (*Solyc03g058160*) in tomato, whose role in glandular trichome development remains unknown (Figure S6). In contrast, SlHAIR (*Solyc10g078970*) and SlZFP8L/SlHAIR2 (*Solyc10g078990*), paralogs of SlZFP8 (Figure S6), are well-characterized C2H2 zinc finger proteins that directly interact with SlWOOLLY to regulate trichome initiation and elongation via SlZFP6 (Zheng et al., 2021, Chang et al., 2018). It was observed that *NtZFP8* is expressed in all types of developing trichomes suggesting a role in the development of these specialized micro-organs (Figure 3A-B). To further investigate the role of NtZFP8 in trichome development, transgenic lines overexpressing *NtZFP8* were generated (Figure 3A). In p*35S::NtZFP8* lines, a significant decrease in trichome density— including both glandular and non-glandular types—was observed(Figure 3C-D). This finding suggests that NtZFP8 might function as a repressor of trichome development, playing an antagonistic role to MIXTA-like transcription factor. Given the contrasting phenotypes of NtZFP8 and AmMIXTA overexpression lines, it is possible that NtZFP8 is induced in MIXTA-induced transgenic lines, where it could modulate AmMIXTA transcriptional activity as part of a negative feedback mechanism to fine-tune trichome development in these lines. Alternatively, the inhibitory effect of NtZFP8 overexpression in p*35S::NtZFP8* lines on trichome development could result from an artificial titration effect, in which NtZFP8 binds and sequesters an endogenous MIXTA-like partner with trichome-promoting activity, acting as a dominant-negative allele, rather than reflecting its normal regulatory function.

To date, evidence for zinc finger proteins acting as negative regulators of trichome development remains limited (Han et al., 2021). While in tomato, SlHAIR2 is considered as a positive regulator of trichome development (Zheng et al., 2021), its KO mutant exhibits an intermediate phenotype, with a reduction in type I and II trichomes but an increase in types III and IV, suggesting that SlHAIR2 may promote the formation of certain trichome types while repressing the development of others (Gasparini et al., 2023). In root hair development, a process sharing common genes with trichome development (Ishida et al., 2008), AtZP1 was identified as a zinc finger protein that negatively regulates root hair development and is transcriptionally controlled by AtGL2 (Han et al., 2020). Further research is therefore needed to clarify the precise mode of action of NtZFP8 in *planta*.

Interestingly, AmMIXTA physically interacts with NtZFP8 in yeast two-hybrid assays (Figure 4). This interaction suggests a possible regulatory pathway linking a MIXTA-like transcription factor and NtZFP8 in trichome development. AmMIXTA does not directly regulate *NtZFP8* expression, as evidenced by DAP-seq results (Table S5). AmMIXTA and NtZFP8 significantly overlap in their directly regulated genes, with 45% of AmMIXTA targets also targeted by NtZFP8 and 77% of NtZFP8 targets also bound by AmMIXTA, indicating their joint or possibly antagonist role as transcriptional regulators. AmMIXTA was shown to bind to the GGTGGTGGTGG motif while NtZFP8 was found to bind the CTTCTTCTTCT motif (Figure 5E-F).

Common targets bound in the promoter region identified by DAP-seq include *NtMYB6* and *NtGLABRA2* (Figure 5D), whose homologs were characterized as involved in trichome development. For instance, *CsMYB6* is a negative regulator of trichome initiation in cucumber (Yang et al., 2018) while *AtGL2* and *CsGL2* promote trichome formation (Rerie et al., 1994, Yang et al., 2011b). Interestingly, SlGL2 is induced by *SlSHINE3*, which also regulates epidermal patterning and cuticle formation in tomato (Shi et al., 2013). In cucumber, CsGL3 (a HD-ZIP IV class protein) and CsGL1 (a HD-ZIP I class protein) interact to initiate multicellular trichome development, activating three regulators: CsGL2, CsMYB6, and CsWIN1/SHINE1 (Liu et al., 2016, Han et al., 2022). Their close tobacco homologs are direct targets of AmMIXTA and/or NtZFP8 (Figure 5D). Co-regulation of genes such as *NtWIN1* by NtZFP8, *NtMYB6* and *NtGLABRA2* by AmMIXTA and NtZFP8, along other known transcription factors and signaling molecules as part of their regulatory network, suggests that at least part of the transcriptional control regulating trichome development is evolutionarily conserved between species belonging to different plant families.

We also advise caution against an overly simplistic view that each plant species has its own distinct transcriptional network governing the development of different trichome types. Our data highlight the fact that some components, like key regulatory modules are actually shared across species. This applies to both glandular and non-glandular trichomes, with genes like ZFP8, GL2, MYB6, WOOLLY/PDF2 playing a role in regulating these processes in species as diverse as *N. tabacum, S. Lycopersicum, C. sativus* or *A. thaliana*. This highlights the need to avoid assuming that all differences in trichome types come from distinct, independent regulatory networks. Instead, certain genetic elements are shared and reused in evolutionary adaptations, while others may evolve specifically in certain species.

In summary, our research leveraged the role of *AmMIXTA* to investigate the underlying transcriptional network controlling long-stalked glandular trichome formation. It is likely that there is an endogenous yet-to-be identified MIXTA-like interacts with NtZFP8 to regulate this process. By partially revealing the complex transcriptional network behind this specific developmental mechanism, our findings lay the foundation for future studies aimed at identifying key genes and transcription factors driving or repressing trichome development in *Solanaceae* species. These investigations will explore signaling pathways, environmental influences, and the potential for targeted gene editing to optimize glandular trichome development, whether through induction or repression of this developmental process. This research could lead to practical applications in plant breeding programs that capitalize on these benefits for agricultural advancements.

## 4. Material and methods

### 4.1 Genetic Constructs

The AmMIXTA coding sequence was provided by Prof. Beverley Glover (University of Cambridge, UK). This sequence was amplified (Table S7), cloned into a pENTR/D-TOPO entry vector (Invitrogen, 45-0218) and was transferred to a pMDC7 destination vector containing a β-estradiol promoter (Curtis and Grossniklaus, 2003) using Gateway Technology (Life Sciences). The NtZFP8 (LOC107828820) coding sequence was amplified by PCR from cDNA generated based on leaf primordia mRNA (Table S7), and recombined into a pDONR221 vector to create the pENTRY L1-NtZFP8-L2. p35S::NtZFP8 was produced by LR Gateway reaction recombining the corresponding entry clone into the pH7WG2D (Karimi et al., 2002). A 1982 kb fragment upstream of the NtZFP8 coding sequence was cloned into pDONR L4-L1r to produce pENTRY L4-pNtZFP8-R1. pZFP8::nlsGFP-GUS was generated by recombining pENTRY L4-pNtZFP8-R1 with pENTRY L1-nlsGFPGUS-L2 into pH7m34GW (Berhin et al., 2019, Karimi et al., 2002). For yeast two-hybrid assays, coding sequences of NtZFP8, AmMIXTA, and nls-GFP were PCR-amplified using primers with BsaI site overhangs (Table S7) and cloned into CK011 and CK012, containing respectively the GAL4 activation and binding domain, under the ADH1 promoter (Gantner et al., 2018), using Golden Gate cloning technology (Kirchmaier et al., 2013). For protein expression in DAP-seq, pENTRY L1-NtZFP8-L2, pENTRY L1-AmMIXTA-L2, and pENTRY L1-nlsGFP-L2 were transferred to the pIX-HALO plasmid via an LR Gateway reaction, resulting in pIX::HALO-NtZFP8, pIX::HALO-AmMIXTA, and pIX::HALO-nlsGFP constructs (Bartlett et al., 2017). For recombinant protein expression, the coding sequence of both genes were amplified (Table S7) and cloned into pQE60 (Qiagen) using Mlu1 and BamHI restriction sites.

### 4.2 Plant Material and Growth Conditions

*N. tabacum cv Petit Havana* (SR1) was used for stable plant transformation. Seeds were surface-sterilized with chlorine gas, sown on half-strength Murashige and Skoog agar plates (MP Biochemicals, 92610024), stratified at 4°C for 3 days, and subsequently transferred to a growth chamber (16-hour light/8-hour dark, 50 μmol photon/m^2^/s, 25 ± 2°C). Seedlings from in vitro cultures were then transferred to Jiffy pots for a week or germinated in Jiffy pots for two weeks before moving to pots with soil in a phytotron (25°C, 16-hour photoperiod, 200 μmol photon/m²/s) for propagation and phenotyping. All constructs in destination vectors were introduced into *Agrobacterium tumefaciens* strain GV3101 and transformed into tobacco using a leaf disk method modified from Horsh (1985). Transgenic lines carrying pEST::AmMIXTA, p35S::NtZFP8 and pNtZFP8::nlsGFP-GUS were generated. Experiments with AmMIXTA were performed on T2 generation while the ones with ZFP8 were conducted on T1.

### 4.3 mRNA Extraction

For the RNA-seq, aerial parts of 15 seedlings were harvested per sample per condition. To study *NtZFP8* expression, various tissues from 6-week-old plants were used, including leaf primordia (<1.5 cm), young leaves (+-10 cm), mature leaves (+-20cm), and mature trichomes. For each of the harvested tissues, tissues from three plants were mixed to form a sample and four samples were harvested to have four biological replicates. For the overexpressing lines of *NtZFP8*, 3 disks from the same leaf used for trichome density were harvested from three independent plants. RNA was extracted using the Spectrum Plant Total RNA Kit with on-column DNase I digestion.1 µg RNA was reverse transcribed and analyzed by RT-qPCR with specific primers (Table S7). Normalization was done by geometric averaging of NtEF2α, NtACTIN and NtUBC2 (Vandesompele et al., 2002). Relative expression levels were calculated using 2^-ΔΔCT^ method. (Schmittgen and Livak, 2008).

### 4.4 RNA-sequencing

Seedlings from independent T1 lines expressing AmMIXTA were screened for trichome proliferation by germinating them on _1/2M_S medium, with 10 μM β-estradiol and checking *AmMIXTA* expression level via RT-qPCR. Selected T2 seeds were grown under control and β-estradiol conditions for 7 days. Aerial parts from 15 seedlings were collected at 4 and 48 hours post-induction for mRNA extraction (see above).

RNA-seq libraries were generated and sequenced by Genewiz (Azenta life science) using Illumina NovaSeq sequencing plateform, yielding at least of 38 million 150 bp paired-end reads per sample (Table S1). Reads were cleaned, trimmed, and mapped to the *N. tabacum* TN90 reference genome (Sierro et al., 2014) using CLC Genomic Workbench. Differentially expressed genes (DEGs) were identified based on normalized counts per million (CPM), with a fold change >1.5 (upregulated genes) or <0.67 (downregulated genes) and a p-value <0.05, using EdgeR in RStudio (Law et al., 2016).

DEGs were analyzed for enriched Gene Ontology (GO) terms using PANTHER classification system (Figure 3) (Mi et al., 2019, Thomas et al., 2022) and visualized with rrvgo (Figure S3) (Sayols, 2023).

### 4.5 DNA Affinity Purification Sequencing (DAP-seq)

DNA from three 6-week-old plant leaf primordia was extracted using the Wizard Genomic DNA Purification Kit (Promega). The DAPseq experiment followed Bartlett et al. (2017)’s protocol. HALO-AmMIXTA, HALO-NtZFP8 and HALO-nlsGFP (negative control) were each expressed three times using the TnT® Coupled Wheat Germ Extract System (Promega, L41030). Index and adaptor were added to the DNA fragments using NEBNext® Multiplex Oligos for Illumina® primers (NEB, E6440S). DNA-seq libraries were sequenced by Macrogen using Illumina NovaSeq 6000 platform yielding at least of 50 million 150 bp paired-end reads (Table S4). Reads were cleaned and mapped (See RNA-seq). Peak scores (>2) and p-values (<0.05) were calculated using the Transcription Factor CHIP-seq tool from CLC Genomic Workbench comparing our samples to the negative control (DNA bound to nlsGFP). Peaks of interest were selected based on their proximity to a gene: 3kb upstream of a START codon and 1kb downstream of a STOP codon.

### 4.6 Microscopy and Phenotyping

Trichomes were observed using an Observer Z1 microscope. Reporter lines were studied using Zeiss LSM710 and Stellaris 8 Falcon confocal microscopes. Image were processed with Zen and ImageJ softwares.

### 4.7 Yeast Two-Hybrid Assay

The yeast two-hybrid assay followed Paiano et al. (2019), using the *pJ696* yeast strain (Lafuente et al., 2000) and nlsGFP as a negative control.

### 4.8 Electrophoretic Mobility Shift Assay (EMSA)

Recombinant proteins were expressed in *E. coli* BL21(DE3) cells, which were transformed with expression vectors. Cultures were grown at 30°C to OD600 0.3–0.5, then induced with 0.5 mM isopropyl β-D-1-thiogalactopyranoside (IPTG) for 5 hours. Cells were harvested by centrifugation, frozen, and lysed in a buffer (50 mM Tris-HCl, pH 8, 500 mM NaCl, 30 mM imidazole, 0.5 mM DTT, 1 mM PMSF, lysozyme, and protease inhibitors) by sonication. The lysate was clarified by centrifugation, and the protein was purified using a Ni-NTA column. After washing, proteins were eluted with 300 mM imidazole. Protein concentration was measured using a Qubit fluorometer. For the EMSA assay, complementary oligonucleotides encoding the MEME motif were synthesized, with one oligo tagged with Cystein3 (Table S7). Equimolar concentrations (10 μM) of oligos were mixed in STE buffer (100 mM NaCl, 10 mM Tris-HCl, pH 8, 1 mM EDTA) and annealed using a PCR thermocycler with the following program: 98°C for 3 minutes, 75°C for 1 hour, 65°C for 1 hour, 37°C for 30 minutes, and 25°C for 10 minutes. For binding assays, the double-stranded DNA substrate (1 nM) was incubated with the purified protein in a binding buffer containing 25 mM Tris-HCl (pH 7.5), 9% glycerol, 75 mM NaCl, 5 mM DTT, 5 mM MgCl_2,_ and 1 mg/mL BSA at room temperature for 20 minutes. For negative controls without proteins, SEC buffer (50 mM Tris-HCl, pH 8, 10% glycerol, 50 mM NaCl, 5 mM Na_2E_DTA) was used. The reaction was separated on a 10% polyacrylamide gel and visualized using a Typhoon scanner (Cytiva).

## 5. Statement and declaration

## 5.1. Acknowledgments

We extend our sincerest gratitude to Prof. Beverley Glover for generously providing the cDNA of *AmMIXT*A, which was crucial for the success of our experiments. We are also deeply thankful to Prof. Michel Ghislain for providing the yeast-2-hybrid system used in this study, to Prof. Jo Ecker for supplying the genetic material needed for the DAP-seq assay and to Louis Gueuning for assistance in the cloning of the EMSA constructs. We deeply appreciate their support and the resources they provided. We are grateful to Louis Gueuning and Gabriel Walckiers for their critical reading and comments on previous versions of this manuscript.

## 5.2 Funding

This work was supported by the Belgian National Fund for Scientific Research (PDR/PGY grant #T.0116.20 to AB, MP and CH, Charge de recherche #FC46171 to AB and FNRS Scientific Impulse Mandate # F.4522.17 to AR and MV), the Swiss National Science Foundation (FNS; Grant#P2LAP3_191271 to AB). MV benefitted from funding from the UCLouvain Special Research Funds (UCLouvain FSR 2017).

### 5.2.1 Author Contributions

AB conducted the foundational experiments for this study and drafted the manuscript. AR, MV and MP were responsible for the creation of transgenic lines and the acquisition of preliminary transcriptomic data, AR participated in the drafting of a preliminary version of this manuscript. MP assisted AB to develop and execute the Yeast Two-Hybrid (Y2H) experiments. BEA and SL offered essential support in analyzing the RNA-seq and DAP-seq data. CH conceived and supervised the whole project and was actively involved in data analysis and writing of the manuscript.

### 5.3 Data availability

RNA-seq and DAP-seq data sets are MIAME compliant and were submitted to the Gene Expression Omnibus (GEO) database under the GEO Series accession number GSE263485 and GSE263486.

