## Supplemental Figures for "Insight into the genetic network governing long-stalked glandular trichome development in *Nicotiana tabacum*"

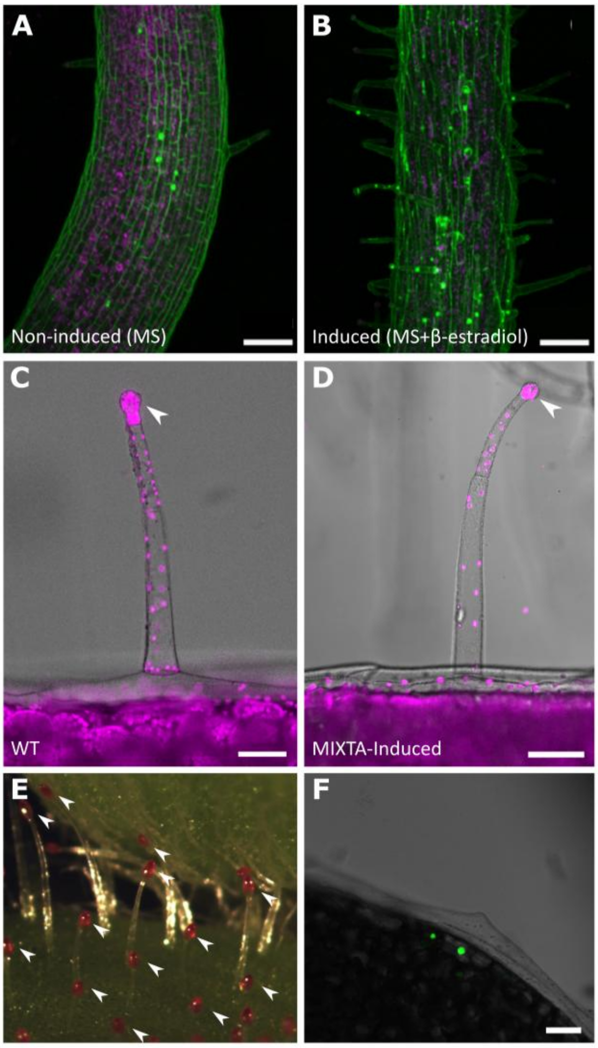


**Figure S1: Detailed phenotypic differences in pEST::AmMIXTA transgenic plants.** Ten-day-old *pEST::AmMIXTA* seedlings were germinated on non-inducing medium (1/2 MS medium) (A) and inducing medium (1/2 MS medium supplemented with 5 µM β-estradiol) (B). Panels (A/B) show the hypocotyls. Magenta represents autofluorescence; green indicates propidium iodide staining. Scale bar: 100 µm.

Panels (C/D) illustrate chlorophyll autofluorescence in the trichome heads of (C) a leaf from a wild-type (WT) plant and (D) a cotyledon from a ten-day-old induced *pEST::AmMIXTA* seedling. White arrowheads point to chlorophyll autofluorescence in the glandular heads. Scale bar: 50 µm. Panel (E) shows induced ten-day-old pEST::AmMIXTA seedlings stained with Rhodamine B, an acylsugar dye that stains the trichome heads red, confirming the full development of trichomes in the induced line. White arrowheads point to stained trichome heads. In panel (F), the nuclear localization of VENUS-tagged MIXTA under the same inducible promoter in inducible conditions is displayed. VENUS-tagged MIXTA is shown in green. The scale bar is 50 µm.

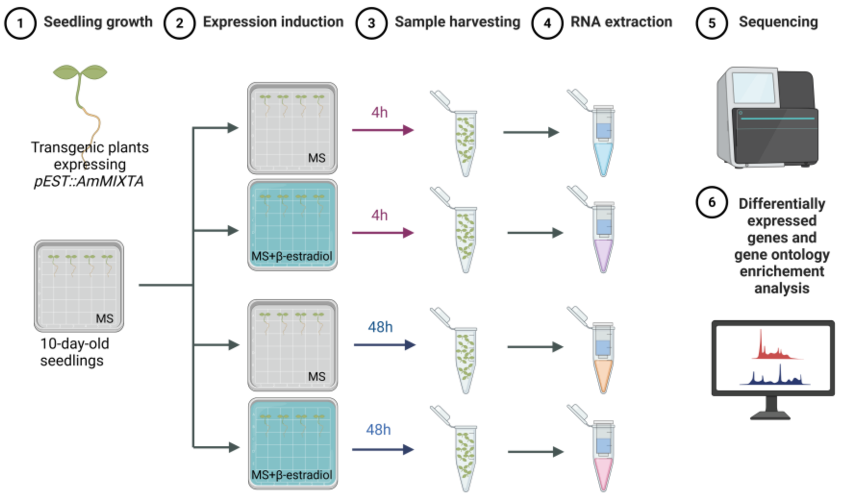


**Figure S2: Utilization of AmMIXTA as a Tool for Identifying Genes Involved in the Early Stages of Trichome Development: Experimental Scheme.** This figure outlines the step-by-step experimental approach employed to elucidate the role of AmMIXTA in the initiation and early development of trichomes. Initially, pEST::AmMIXTA transgenic plants were cultivated under inducing conditions to induce the expression of AmMIXTA, the same line was grown in non-inducing conditions as a control. Subsequent stages involved the collection of samples at critical early developmental stages, followed by RNA extraction and high-throughput sequencing to identify differentially expressed genes. The schematic representation includes key experimental steps such as treatment conditions, sample collection time points, and the analytical methods used to dissect the genetic pathways activated by AmMIXTA in trichome development. Created in BioRender. Berhin, A. (2025) https://BioRender.com/68t0n30.

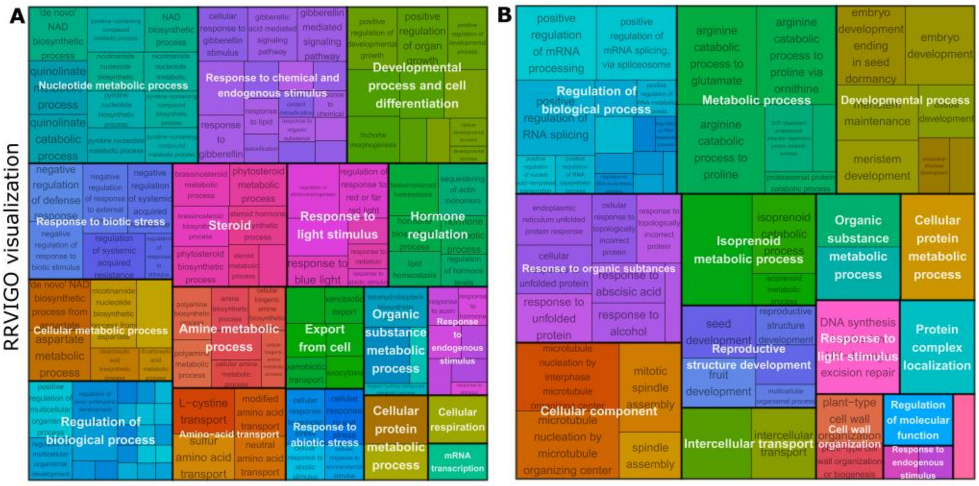


**Figure S3: GO Term Enrichment Visualization with Rrvigo**

Utilizing Rrvigo (Sayols, 2023), a tool designed to visualize enriched GO terms, upregulated GO terms identified in Figure 3 at 4 hours (A) and 48 hours (B) following pEST::AmMIXTA induction were categorized into subgroups. This visualization facilitates the understanding of the functional significance of genes upregulated in response to pEST::AmMIXTA induction, by grouping similar GO terms together and highlighting the primary biological processes involved.

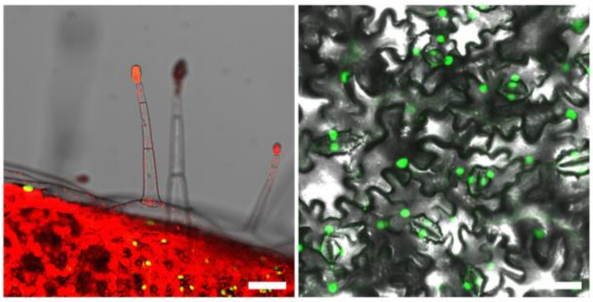

**Figure S4: NtZFP8 Expression Pattern in 6-week-old leaves.**

Transgenic plants expressing pZFP8::nlsGFP-GUS in larger leaves (15 cm) of 6 week-old plants. Left panel shows a trichome (note that mature glandular trichomes are not labelled); right panel shows the epidermal cell layer. Scale bars represent 100 μm; green indicates nlsGFP; red denotes chlorophyll autofluorescence.

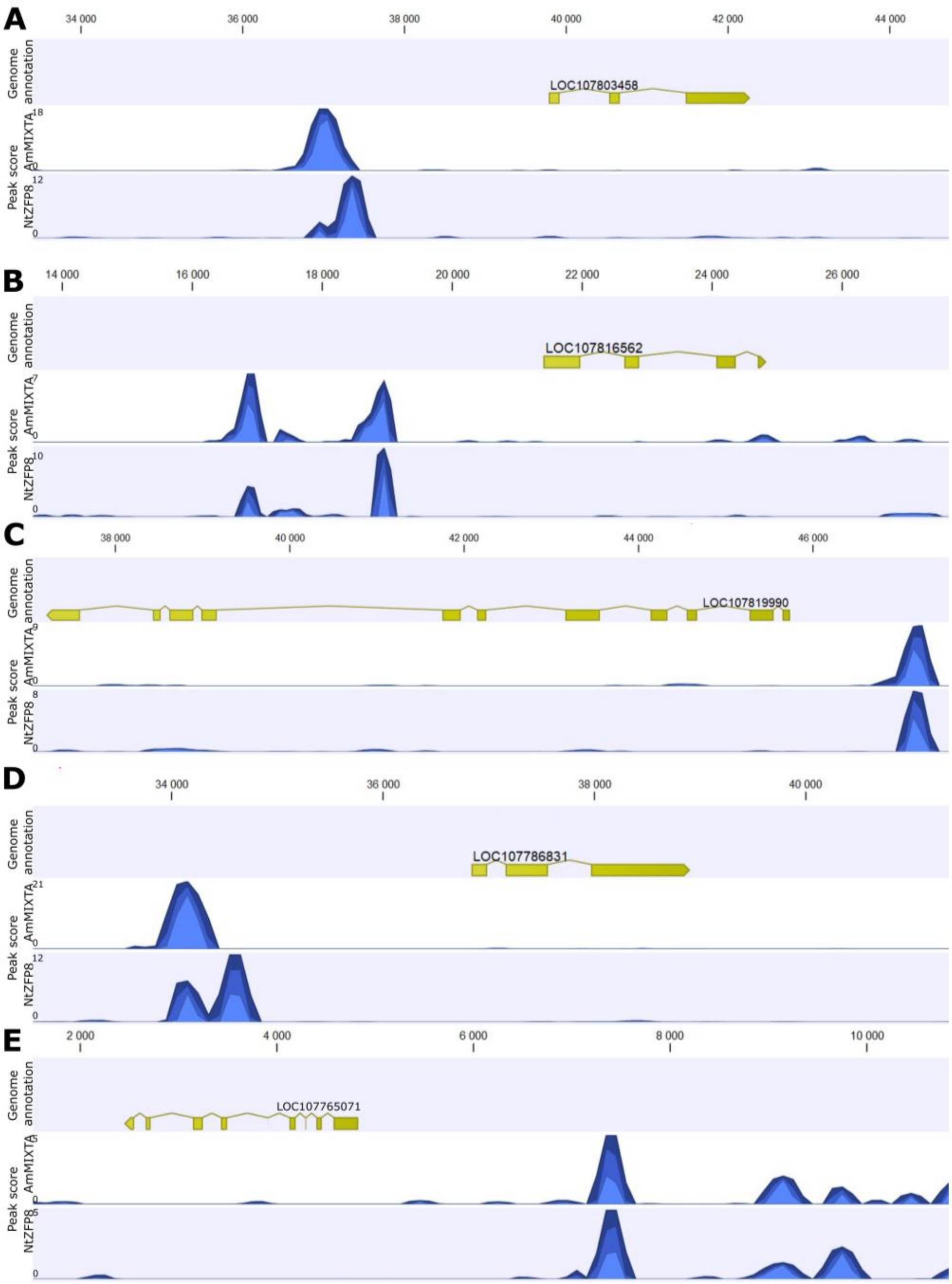


**Figure S5: Top 5 Binding Sites of NtZFP8 and AmMIXTA Identified by DAP-seq.** DAP-seq analysis identified genes whose promoters are targeted by both AmMIXTA and NtZFP8: (A) LOC107803458, NtMYB6; (B) LOC107816562, NtCONSTANS-like 10; (C) LOC107819990, NtGLABRA2; (D) LOC107786831, NtJACKDAW; and (E) LOC107765071, NtAGL66. In each panel, the upper track displays the gene annotation and its position on the contig, while the lower track shows the DAP-seq peak shape score and the peak's localization. The analysis and creation of this figure were performed using CLC Genomics

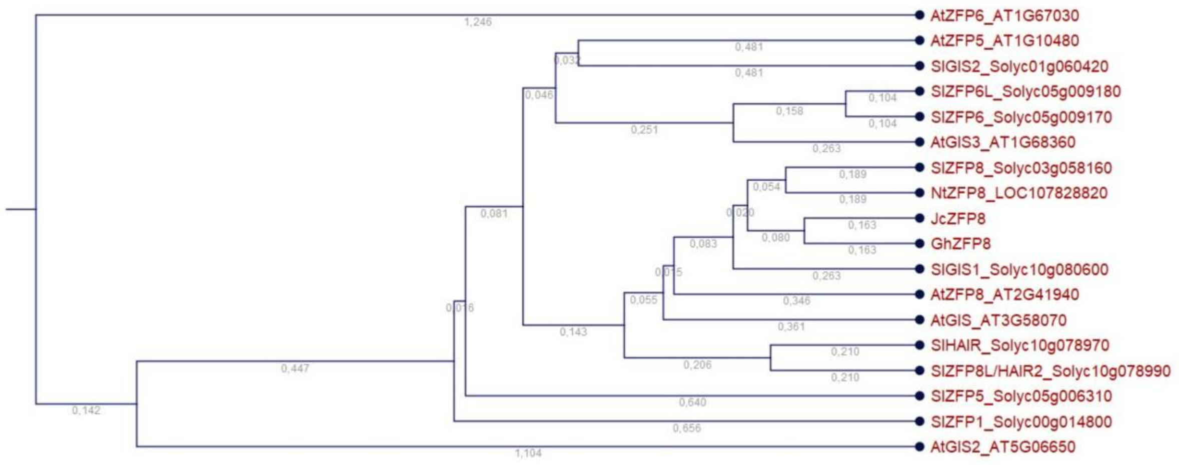


**Figure S6: Phylogenic tree of ZINC-FINGER proteins.** Phylogenic relationship of ZINC-FINGER proteins in tomato, Arabidopsis, coton and *Jatropha curcas*. Bootstrap analysis was based on 1,000 replicates (CLC workbench, QIAGEN). The number of base pair substitutions/sites is indicated at each branch.

**Supplemental Table S1: Exonic read mapping to the *N.tabacum* genome before and after trimming steps**

**Supplemental Table S2: Upregulated DEGs at 4 h and 48 h post-induction of *AmMIXTA***

**Supplemental Table S3: GO term enrichment analysis at 4 h and 48 h post-induction of *AmMIXTA***

**Supplemental Table S4: DAP-seq Read mapping to the *N.tabacum* genome before and after trimming steps**

**Supplemental Table S5: DAP-seq table for AmMIXTA binding sites**

**Supplemental Table S6: DAP-seq table for NtZFP8 binding sites**

**Supplemental Table S7: PCR primers used in this study**
